# The Cystic Fibrosis Transmembrane Conductance Regulator (CFTR) Modulates the Functional Output of Human Taste Bud Cells

**DOI:** 10.64898/2025.12.31.697221

**Authors:** Satya Iyer, Jean-Pierre Montmayeur, William D. Hunt, Cedrick D. Dotson

**Affiliations:** Neuroscience Institute, Georgia State University, Atlanta, GA, USA; School of Electrical and Computer Engineering, Georgia Institute of Technology, Atlanta, GA, USA

## Abstract

We recently discovered that the cystic fibrosis transmembrane conductance regulator protein (CFTR), which functions as a channel that transports chloride and bicarbonate across epithelial surfaces, is expressed in both human and murine TBCs, but how it functions in these cells remains unknown. We postulated that CFTR may impact peripheral taste signaling at the level of taste receptor-expressing cells of the taste bud. To begin to test this hypothesis, we assessed how pharmacological manipulation of CFTR could affect the functional responses of human fungiform taste bud cells to prototypical taste stimuli (e.g., bitter, sweet, fat) using single cell calcium imaging and neurotransmitter (ATP) release measurements. We first established the presence of CFTR in these cells using immunocytochemistry and RT-PCR. We next found that CFTR inhibition generally increased stimulus-evoked calcium responses but that the specific response parameters impacted varied across different stimuli, likely due to differences in signal transduction mechanisms and the involvement of store-operated calcium channels. For example, response amplitudes to bitter and sweet stimuli were significantly enhanced with no changes in the proportion of cells responding to these stimuli whereas the opposite trends were observed with a fatty acid stimulus. Additionally, bitter-evoked neurotransmitter release was significantly enhanced by CFTR inhibition, suggesting that this effect is reflected throughout the signal transduction cascade. Ongoing and future experiments are utilizing shRNA knockdown as well as intracellular and extracellular electrophysiology to further interrogate the impacts of CFTR. In addition to human TBCs, we have detected CFTR in mouse taste tissues. Moreover, we have mined mouse TBC RNA sequencing datasets to determine CFTR co-expression patterns to inform future cellular and behavioral experiments in mice. Taken together, these data suggest that CFTR can modulate the signaling output of the taste bud.

## INTRODUCTION

The sense of taste acts to protect the rest of the alimentary canal by providing information on which nutrients to ingest and which to reject (e.g., [1–12]), which is critical for an organism’s survival. Taste buds, which are clusters of taste bud cells (TBCs), the primary sensory cells of the gustatory system, express multiple classes of receptors that mediate our perceptions of taste quality (e.g., sweet, bitter, umami, sour). The receptors are largely expressed in non-overlapping sets of different TBC types. In addition to expressing metabotropic and ionotropic taste receptors, TBCs also express several additional types of ion channels; the activity of which is modulated directly or indirectly upon stimulation of the cells with tastants. Such channels, including epithelial sodium channels (ENaCs), otopetrin 1 (OTOP1), transient receptor potential cation channel subfamily M member 4 (TRPM4), and TRPM5, when deleted or inactivated, have profound impacts on TBC physiology and taste perception (e.g., [13–16]).

We recently discovered that the cystic fibrosis transmembrane conductance regulator protein (CFTR), which functions as a channel that transports chloride and bicarbonate across epithelial surfaces, is expressed in both human and murine TBCs. The expression of the CFTR channel has previously been reported in rat TBCs [17]. However, how it functions in these cells, if at all, remains unknown.

We hypothesized that CFTR plays a critical role in TBC physiology and function. To better understand the roles of CFTR throughout the peripheral gustatory system, we first attempted to identify which TBC types express this channel. To do this, we performed expression analyses of CFTR in TBCs together with established TBC markers in both human and murine TBCs. We also mined publicly available murine TBC RNAseq datasets to assess expression patterns using RNA-based methodology. We then asked if the inhibition of CFTR channels leads to changes in the response of TBCs when stimulated by prototypical gustatory stimuli. Approaches used to dissect CFTR’s role in taste signaling included immunohistochemistry, single-cell RT-PCR, RNAScope, calcium imaging, shRNA gene knockdown, ATP secretion assays, and electrophysiology.

We report that CFTR is expressed in a subset of tastant responsive TBCs. We also found that CFTR inhibition generally increased stimulus-evoked calcium responses but that the specific response parameters impacted varied across different stimuli, likely due to differences in signal transduction mechanisms and the involvement of store-operated calcium channels.

Our results could help to provide a greater understanding of how tastants are transduced by TBCs and how CFTR’s function influences the overall output of the taste periphery to the central nervous system. We hope to set the stage for future advancements in the fields of sensory neuroscience, epithelial physiology, and ion channel biology, and reveal generalizable principles of cellular communication.

## METHODS AND MATERIALS

### Cell Culture of Human TBCs

Human fungiform taste bud cells (hTBCs) are SV40 immortalized human fungiform taste bud cells (Applied Biological Materials) and have been previously characterized [18–21]. They were cultured in Prigrow® V Medium (Applied Biological Materials) supplemented with 10% fetal bovine serum (Gibco) and 1% penicillin/streptomycin cocktail (Cytiva) at 37°C in a C170i humidified incubator (Eppendorf) maintained at 5% carbon dioxide in air. These cells were passaged between one and five times after thawing (and never more than ten times total) before experimental assessments. Furthermore, any between-cell assessments were conducted on hTBCs unfrozen from the same batch and passaged the same number of times, give or take one passage.

### Animals

This study was approved by the Institutional Animal Care and Use Committee (IACUC) at Georgia State University. All procedures in the study were carried out in accordance with the principles of the National Research Council’s guide for the care and use of laboratory animals. Male CALHM1-WT with a 129Sv ×C57BL/6J genetic background were a generous gift from Dr. Phillipe Marambaud (Feinstein Institutes for Biomedical Research, NY). All animals were housed at 22-24°C on a 12/12 h light cycle and fed standard chow and water *ad libitum*. Homozygous WT or KO mice were used for breeding and genotypes confirmed every three generations (Transnetyx).

### Experimental Solutions and Reagents

Tyrode’s buffer consisted of 140 mM NaCl, 5 mM KCl, 10 mM HEPES, 1 mM CaCl_2_, 1 mM MgCl_2_, 10 mM glucose (Alfa Aeser) and 10 mM sodium pyruvate in double distilled water and brought to a pH of 7.4 with NaOH. Artificial saliva consisted of 5 mM NaCl, 5 mM KCl, 6 mM NaHCO_3_, 1.3 mM HCl, 0.25 mM CaCl_2_, 0.25 mM MgCl_2_, 0.12 mM K_2_HPO_4_ and .12 mM KH_2_PO_4_. All chemicals were purchased from Sigma Aldrich unless otherwise specified. Linoleic acid stock solutions were made in 100% ethanol and stored under nitrogen at -20°C. Stock solutions of the CFTR inhibitor CFTR-inh-172 (Sigma-Aldrich Inc., Cat. # 219670) were made in dimethyl sulfoxide and stored at -20°C. All experimental solutions were freshly made in Tyrode’s buffer or artificial saliva on the day of the experiment.

### Mouse Taste Bud Cell (mTBC) Isolation

Mouse circumvallate and fungiform taste cells were isolated from the tongues of male and female CALHM1 WT mice adapting procedures described in [22]. Briefly, 4–8 weeks old mice, were euthanized using carbon dioxide, followed by cervical dislocation. The tongue was removed and placed in a modified Tyrode’s solution composed of 140 mM NaCl, 5 mM KCl, 2 mM CaCl_2,_ 1 mM MgCl_2_, 10 mM HEPES, 10 mM D(+)-glucose, and 1 mM Na pyruvate (pH 8.0). An enzyme solution containing dispase II (2.5 mg/mL - Sigma), and collagenase A (1 mg/mL - Roche) in modified Tyrode’s solution was injected under the dorsal lingual epithelium in the vicinity of the circumvallate papillae or the tip of the tongue where fungiform papillae are located. The tongue was incubated for 25 min at 37°C. Following the incubation, the dorsal lingual epithelium containing the circumvallate and fungiform taste buds was peeled off the underlying muscle layer with forceps, shredded with fine scissors and incubated in 0.25% trypsin in EDTA (Gibco) for 30 min at 37°C to dissociate the tissue. The trypsin was inactivated in Prigrow media supplemented with fetal bovine serum and antibiotics. Large chunks and debris were removed using a 70 μm nylon tissue strainer (Fisher).

### RT-PCR

**Human -** RNA from HuFFs and HEK293 cells were extracted using the RNAeasy mini kit (Qiagen). Total human tongue, lung, and muscle RNA (Biochain) was reverse transcribed using the Maxima H cDNA synthesis kit (Fisher). RT-PCR was performed on 400ng of total RNA using custom designed primers (Supplementary Table 1) designed to cross an intron (Primer-Blast). Conditions Tm: 56°C - 50 cycles (Mastercycler, Eppendorf). Products were separated on a 1.5% agarose gel. Images were captured with a UVP system.

**Mouse** – RNA from dissociated mTBCs was extracted using the RNAeasy mini kit (Qiagen) and reverse transcribed using the Maxima H cDNA synthesis kit (Fisher). RT-PCR was performed on ¼ of the cDNA reaction using custom designed primers (supplementary Table 1) designed to cross an intron (Primer-Blast). Conditions Tm: 56°C - 50 cycles (Mastercycler, Eppendorf). Products were separated on a 1.5% agarose gel. Image were captured with a UVP system.

### CFTR knockdown

HuFF cells were plated at a density of 1.5x10^5^ cells in 6cm dishes containing either the control shRNA lentiviral particles (sc-108080) or the CFTR shRNA lentiviral particles (sc-35054-V, Santa Cruz Biotechnology) and 10 mg/ml of Polybrene (Santa Cruz). After 24-48h of incubation with the lentivirus, cells were trypsinized and allowed to grow in 10-cm^2^ dishes in culture medium supplemented with 0.5 mg/ml of puromycin (Thermo Scientific) to enrich the population in CFTR knockdown cells. When selection dishes reached 30% confluency, they were trypsinized and plated on coated glass coverslips for calcium imaging or 6 cm dish for RNA extraction followed by qRT-PCR to verify the efficiency of the CFTR knockdown.

### Quantitative reverse transcriptase polymerase chain reaction (qRT-PCR)

Total RNA was extracted from HuFF cells using the RNeasy® Plus Mini Kit (Qiagen). RNA was converted to cDNA using the Superscript VILO cDNA Synthesis Kit (Thermo Fisher Scientific). TaqMan® Assays (Roche Molecular Systems, Inc.) were obtained from Thermo Fisher Scientific (see Table S1). Duplex qRT-PCR reactions using two different fluorescent dyes were conducted for each gene of interest with GADPH as a simultaneous internal control. Final reaction mixture (20 µL) contained: 1 µL TaqMan Gene Expression Assay *GAPDH* probe (VIC-labelled) (20 ×), 1 µL TaqMan Gene Expression Assay probe (FAM-labelled) (20 ×), 10 µL TaqMan Gene Expression Master Mix (2 ×), 4 µL cDNA template (∼400 ng) and 4 µL nuclease-free water. Quantitative real-time PCR analyses were conducted on a QuantStudio 3 Real-Time PCR system (Thermo Fisher Scientific) using the following parameters: 2 min hold at 50°C, 10 min hold at 95°C, then 15 sec at 95°C followed by one minute at 60°C for 50 cycles. Relative gene expression was then quantified by using the 2^−ΔΔCT^ analysis method. All cycle threshold (C_T_) values (with a cut-off at >40 cycles) were averaged across replicates, normalized to *GAPDH* to produce ΔC_T_ values, then further normalized to the ΔC_T_ of the calibrator gene *PLCβ2*, which was quasi-arbitrarily selected given its common use as a TBC marker, to generate ΔΔC_T_ values. Relative gene expression was determined by resolving 2^−ΔΔCT^ for each independent experiment of each gene of interest. At least three independent experiments consisting of four replicates each were conducted for each gene of interest.

### Immunohistochemistry

Mouse circumvallate papillae were isolated from the tongues of male CALHM1 WT mice adapting procedures described in ([22]). Briefly, 4–8 weeks old mice, were euthanized using carbon dioxide, followed by cervical dislocation. The tongue was removed and immersed in 4% paraformaldehyde/0.1M PBS pH7.4 overnight at 4°C followed by overnight incubation in 30% sucrose/0.1M PBS, pH 7.5 prior to embedding in OCT compound (Fisher). 12μm cryosections were rehydrated in PBS pH7.4 10 min at RT. Antigen retrieval was performed in 10 mM sodium citrate, pH 8.5 at 80°C for 10 min. Sections were then washed 3 times 5 min in PBS-0.1% Triton X-100 prior to incubation in blocking solution (10% donkey serum in PBS-0.1% Triton X-100) for 30 min at 25°C. Primary antibody was added to the sections in blocking solution and incubated overnight at 4°C (Rabbit anti-CFTR (ACL006, Alomone, RRID:AB_2039804, 1/500) and Goat anti-Gustducin (OAEB00418, Aviva System Biology Corp., RRID:AB_10882823, 1/500); sections were washed in PBS-0.1% Triton X-100 3 times 10 min at RT and then incubated with the appropriate secondary antibody (Donkey anti-Rabbit Alexa Fluor 546, Fisher Scientific, A10040 RRID: AB_2534016, (1/500) or Donkey anti-Goat Alexa Fluor 488, Fisher Scientific, A11055 RRID:AB_2534102, (1/500)) at RT for 2h in the dark. Controls with no primary antibody were included in each experiment. After secondary antibody incubation, sections were washed in PBS (3x10 min) and mounted using Fluoromount G (Southern Biotechnology Associates, Birmingham, AL) containing DAPI (BD Pahrmingen) and coverslipped. All images were obtained using a Zeiss LSM 700 Confocal Microscope (Zeiss, Oberkochen, Germany). Stacks were collected using Zen software (Zeiss) and images were processed using Adobe Photoshop CS 2025 software adjusting brightness and contrast.

### Immunocytochemistry

hTBCs were seeded onto CC2-treated chamber slides (Lab Tek). 24-48 hours post-seeding cells were fixed with 4% paraformaldehyde (PFA; w/v) for 20 min at room temperature. Subsequently, the cells were permeabilized in 0.1M PBS pH 7.4, 0.5% Triton X-100 for 10 min twice. Then, the cells were incubated in 10mM sodium citrate pH6.0 10 min at 85°C and rinsed twice for min with 0.1M PBS pH 7.4, 0.1% Triton X-100 prior to incubation in blocking solution (10% blocking agent – Roche - in 0.1M PBS pH 7.4, 0.1% triton x-100) for 30 min at room temperature. The Primary antibodies (1:500 rabbit polyclonal anti-NPY1R, Bioss, (bs-1070R; RRID:AB_10856532) and 1:100 rabbit polyclonal anti-NPY2R, Bioss (bs-0937; RRID:AB_10856382) were diluted in blocking solution and applied onto the cells for incubation overnight at 4°C. After three rinses of 10 min each in 0.1% Triton X-100 in PBS pH 7.4 the cells were incubated with 1:500 diluted donkey-anti rabbit Alexa Fluor 488 (A-21206; RRID:AB_2535792) or 594 (A-21207; RRID:AB_141637) (Invitrogen) for 90 min at room temperature in the dark. Then the cells were washed twice in PBS pH 7.4 at room temperature and nuclei stained with DAPI (BD Pharmingen) in PBS pH 7.4 for 5 min at room temperature. The sections were rinsed again, twice in PBS pH 7.4 and coverslips were mounted with Fluoromount G (Southern Biotech). Negative control experiments without primary antibody and/or with blocking peptides (when available) were performed to assess specificity of the primary & secondary antibodies (i.e., background signal generated by the fluorescent secondary antibodies). Images were collected with an ECHO fluorescent microscope (ECHO) and processed with the ECHO proprietary software and/or ImageJ.

### Live Cell Calcium Imaging

hTBCs were seeded on glass coverslips coated with a rat tail collagen extracellular matrix (Applied Biological Materials) diluted to 10% v/v in phosphate-buffered saline devoid of calcium and magnesium. The cells were allowed to settle overnight in culture medium, then incubated in Tyrode’s buffer containing 2 μM fura 2-AM and 0.05% Pluronic F-127 (Thermofisher Scientific) for 30 min in the dark in the incubator at 37°C. Compounds used for preincubation experiments were also added to the buffer as part of this incubation step. The coverslip was then set on a recording chamber (RC-26, Warner Instruments) with vacuum grease and mounted onto a widefield inverted epifluorescence microscope (IX-83, Olympus Life Science). Cells were kept in Tyrode’s buffer and photoexcited at 340nm and 380nm wavelengths in quick succession using a pe340 LED light source (CoolLED) while emissions at 510nm to both wavelengths were captured with a sCMOS camera (Hamamatsu Photonics) every 3 sec. Drug solutions were applied to the recording chamber for 90-120 sec in a dropwise manner then subsequently washed with clean Tyrode’s buffer for at least 2 min to allow the cells to return to a stable baseline. Raw fluorescence emissions were recorded, and a 340/380 emission ratio was calculated for each timepoint using MetaFluor software (Molecular Devices). This ratio was subsequently used as a relative measure of cytosolic calcium levels.

### Extracellular Adenosine Triphosphate (ATP) Measurements

hTBCs were seeded in tissue culture treated 24-well plates and kept in culture media for 24-48 h. At a confluence of 70%, approximately 10-15,000 cells per well, the cells were washed once and incubated in Tyrode’s buffer in the presence or absence of CFTR-inh-172 for 30 min at 37^◦^C. After 4 min of incubation, two 50 μL aliquots of buffer were removed from each well and placed in a white 96-well plate (Corning, Inc.). This plate was then placed in a H1M multimodal microplate reader (Agilent Technologies) and bioluminescence was measured immediately following injection of a luciferase assay mix (HS Kit II, Roche). The bioluminescence measure was converted to an ATP concentration using a linear regression fit of serially diluted ATP standard solutions. At the conclusion of the incubation period, the Tyrode’s buffer was aspirated from each well and 450 μL of artificial saliva containing stimuli was added to each well. The plate was then incubated at 37°C for four minutes and two 50 μL aliquots per well were removed and assessed for ATP concentrations as described above. For both measurements, concentrations were averaged across both technical replicates. The second concentration for each well was divided by the first to produce an ATP concentration ratio and correct for differences in baseline ATP concentrations across wells.

### Data Analysis

For calcium imaging, a cell was considered responsive if (i) during the introduction of that stimulus it exhibited (ii) a 340/380 ratio increase greater than two standard deviations over baseline (averaged over sixty seconds of initial recording) that (iii) returned to a stable baseline when the stimulus was removed. Response rates were defined using the proportion of cells responding per total cells assessed for each condition. Either a chi square test, McNemar’s chi square test, or Cochran’s Q test were used with Yates continuity corrections where appropriate to determine statistically significant differences in response rates. Response amplitudes were calculated by taking the peak 340/380 ratio of cells responding to stimuli and subtracting the corresponding baseline. All response amplitudes were analyzed using a within-subjects design; cells were stimulated twice with different stimuli, and the second amplitude was normalized to the first amplitude (or *vice versa*, depending on the experimental question) to account for cell-specific differences in dye loading and responsivity. Statistical differences in relative response amplitudes were analyzed using a Wilcoxon signed rank sum test. For ATP measurements, ATP concentration ratios were analyzed using two-way ANOVAs with stimulus and incubation buffer as factors, and Bonferonni-corrected *post hoc* testing was used when applicable. Significant statistical differences for all measures were determined using a cut-off of p < 0.05. RStudio and the R programming language were used to conduct statistical testing and generate figures, with the exception of the representative traces of calcium imaging, which were generated using MetaFluor software and Adobe Illustrator (Adobe).

### Automated Patch Clamp Electrophysiological Recordings

Whole cell patch clamp recordings were conducted using a Port-A-Patch Basic automated patch clamp rig (Nanion Technologies, Inc., Munich, Germany). hTBCs were cultured to a confluence of 70%, detached from the culture dish using TrypLE™ enzyme (Fisher Scientific) for two minutes, then suspended in Prigrow V medium (Applied Biological Materials) under constant shaking until they were used for electrophysiological experiments. An aliquot of the cell suspension was placed on a 3-5 MΩ resistance chip and loaded onto the Port-A-Patch rig. Gentle suction was applied through the chip to bring a cell to the hole that forms the recording electrode. Then, a barium-based seal enhancer (in mM: 80 NaCl, 60 NMDG, 5 KCl, 10 BaCl_2_, 1 MgCl_2_, 5 glucose and 10 HEPES, pH 7.4, 302 mOsm) was introduced to help produce a GΩ seal. Suction was increased in order to perforate the cell and enter a whole-cell configuration. A sulfate-based internal solution (90 Cs_2_SO_4_, 20 CsCl, 10 HEGTA, 10 HEPES, ph 7.2, 285 mOsm) and barium-based external solution (80 NaCl, 60 NMDG, 5 KCl, 5 BaCl_2_, 1 MgCl_2_, 5 glucose and 10 HEPES, pH 7.4, 298 mOsm) were used to isolate chloride currents. Current recordings were performed in voltage clamp with an initial holding potential of -80 mV.

## RESULTS

### The CFTR is expressed in human and mouse TBCs

Mice are a preferred model for CFTR pre-clinical research due to their anatomical, physiological, and genetic similarity to humans and because of the availability of numerous genetically modified strains available for physiological and behavioral studies. To confirm that the CFTR was present in mouse TBCs, my lab first conducted immunohistochemistry (IHC) on mouse circumvallate papillae (CV). Our data confirms that some gustducin (Gust)-positive (+) cells co-express the CFTR (**Fig. 1**) which is consistent with the Yamada *et al.*, Gust expressing RNAseq dataset [23] as well as Merigo *et al.* [17], immunostaining data derived from the CV of rats[17]. RT-PCR experiments conducted in TBCs from mouse circumvallate and fungiform papillae show that the CFTR channel is expressed in these cells (**Fig. 2**).

**Figure 1:**
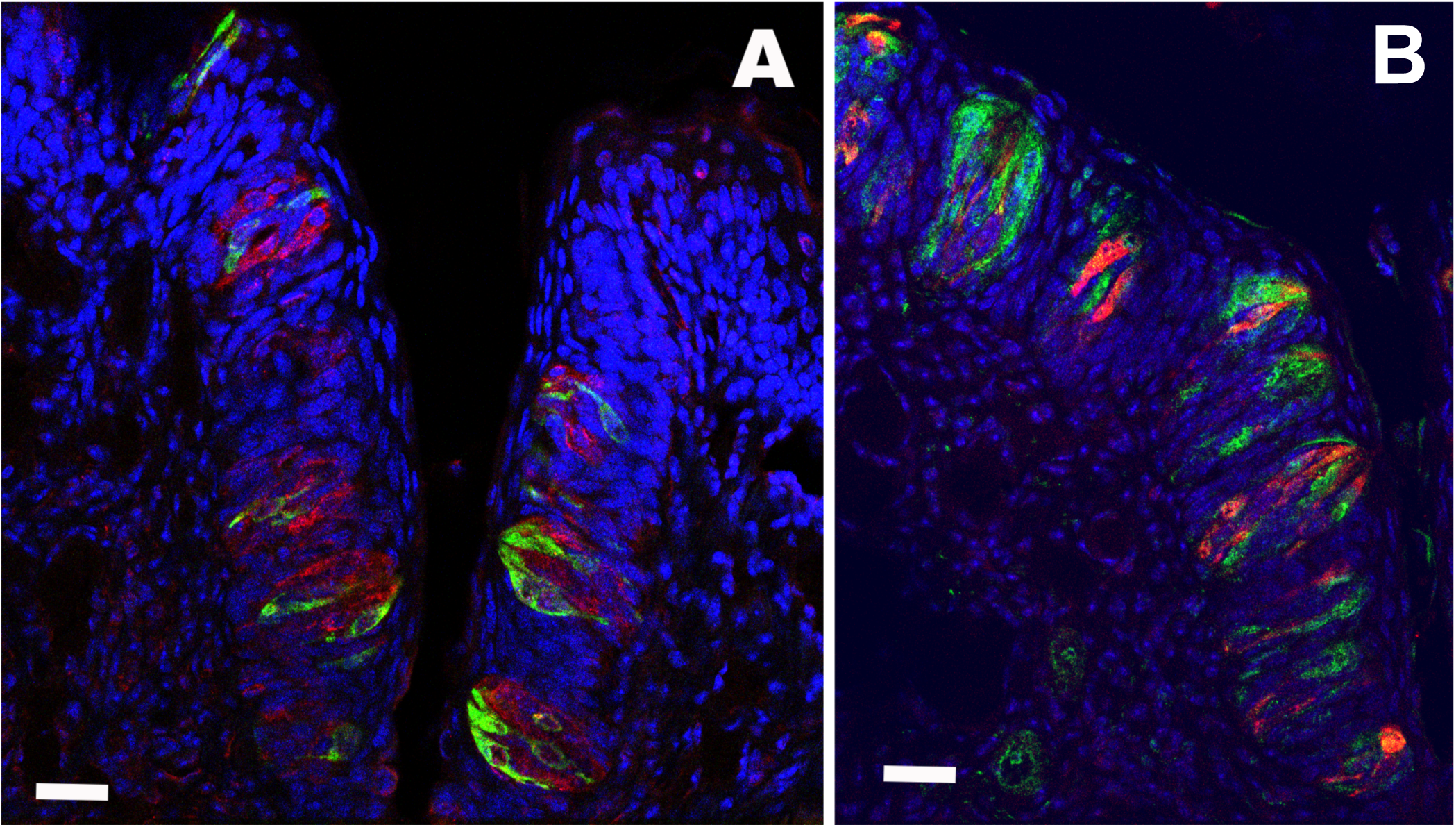
Confocal images of mouse circumvallate (CV) papillae cross-section through the sulcus lined with taste buds. **A-** Shows the localizations of CFTR (red) and gustducin (green), a type 2 taste receptor cell-type marker, in TBCs. **B-** Visualization of CFTR (green) and gustducin (red) in CV papillae section. Secondary antibodies were switched to rule out cross-reactivity and non-specific secondary signal (not shown). Scale bar: 30 μm.

**Figure 2:**
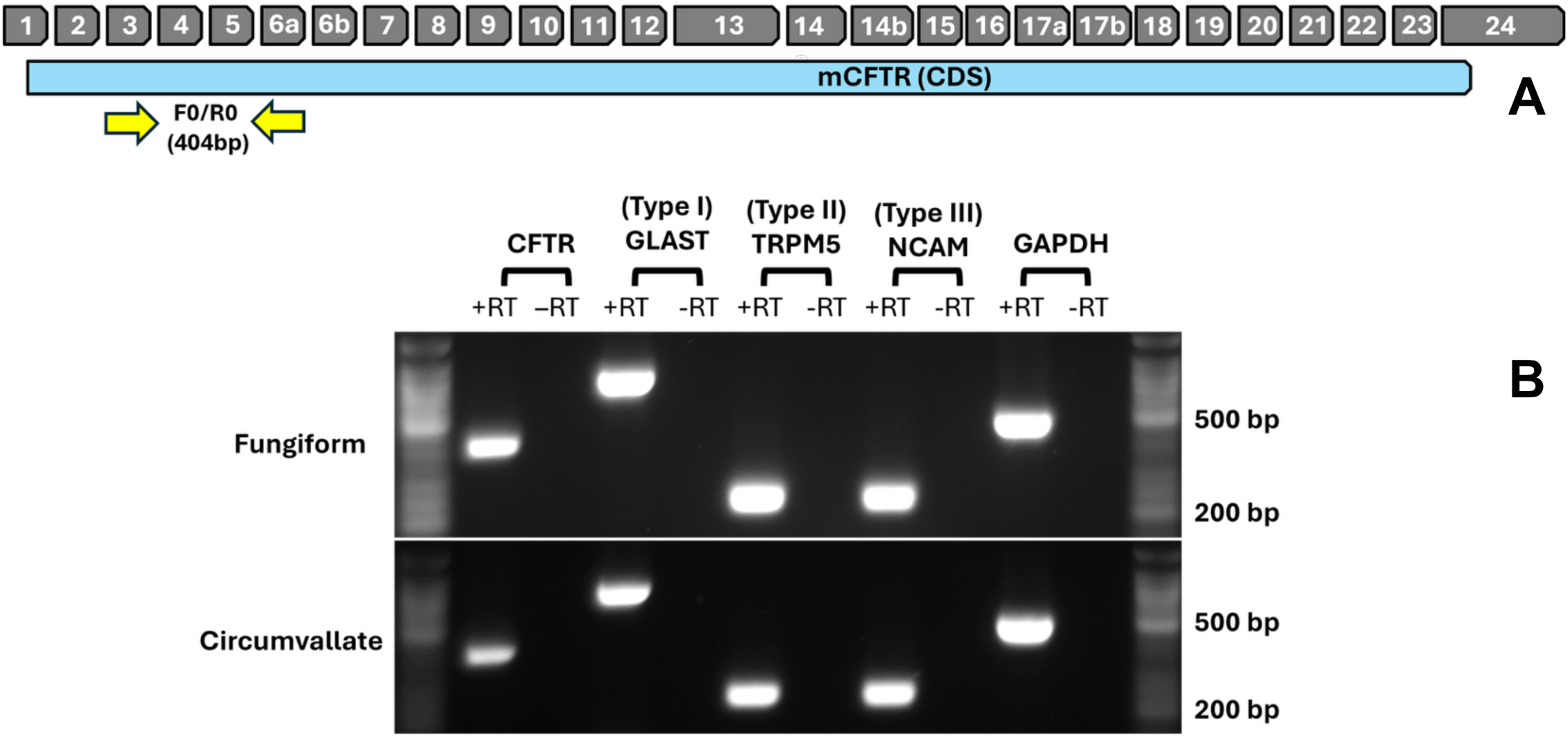
The CFTR transcript is detected in murine TBCs from circumvallate and fungiform papillae. **A-** Mouse CFTR gene structure, coding region, and location of the primers used to amplify a 404 bp fragment across exon 4-5. **B-** Agarose gel showing RT-PCR fragments of CFTR along with TBC cell type specific markers amplified from mouse TBCs from circumvallate and fungiform papillae. No reverse transcriptase (-RT) controls verify the absence of contamination.

We also know from our IHC experiments that, as reported by Merigo *et al.* [17], in rats, the CFTR is also present in some Gust negative TBCs, which we speculate could be type 3 cells (based on the Sukumaran *et al.*, RNAseq dataset [24]; **Table 1**). Indeed, about half (5 of 9) of T1R3 expressing type 2 cells express the CFTR at various levels. A subset (3 of 14) of KCl responsive type 3 cells (∼20%) express the CFTR at various levels. This data is in line with our IHC showing partial overlap in mouse CV co-immunostained for Gust and CFTR and suggests that some CFTR+/Gust-cells may be type 3 cells.

To draw a link with human physiology, we investigated expression of the CFTR in human fungiform TBCs in culture, both by RT-PCR and immunocytochemistry. RT-PCR experiments conducted in these TBCs show that the CFTR channel is expressed in this TBC line (**Fig. 3**). Immunostaining results also confirm the expression of CFTR in human fungiform (HuFF) cells (**Fig. 3c**). HuFF cells, to the best of our knowledge, are the first and only commercially available, immortalized human TBC line (Applied Biological Materials, Richmond, BC, Canada) and have been partially characterized [18–20].

**Figure 3:**
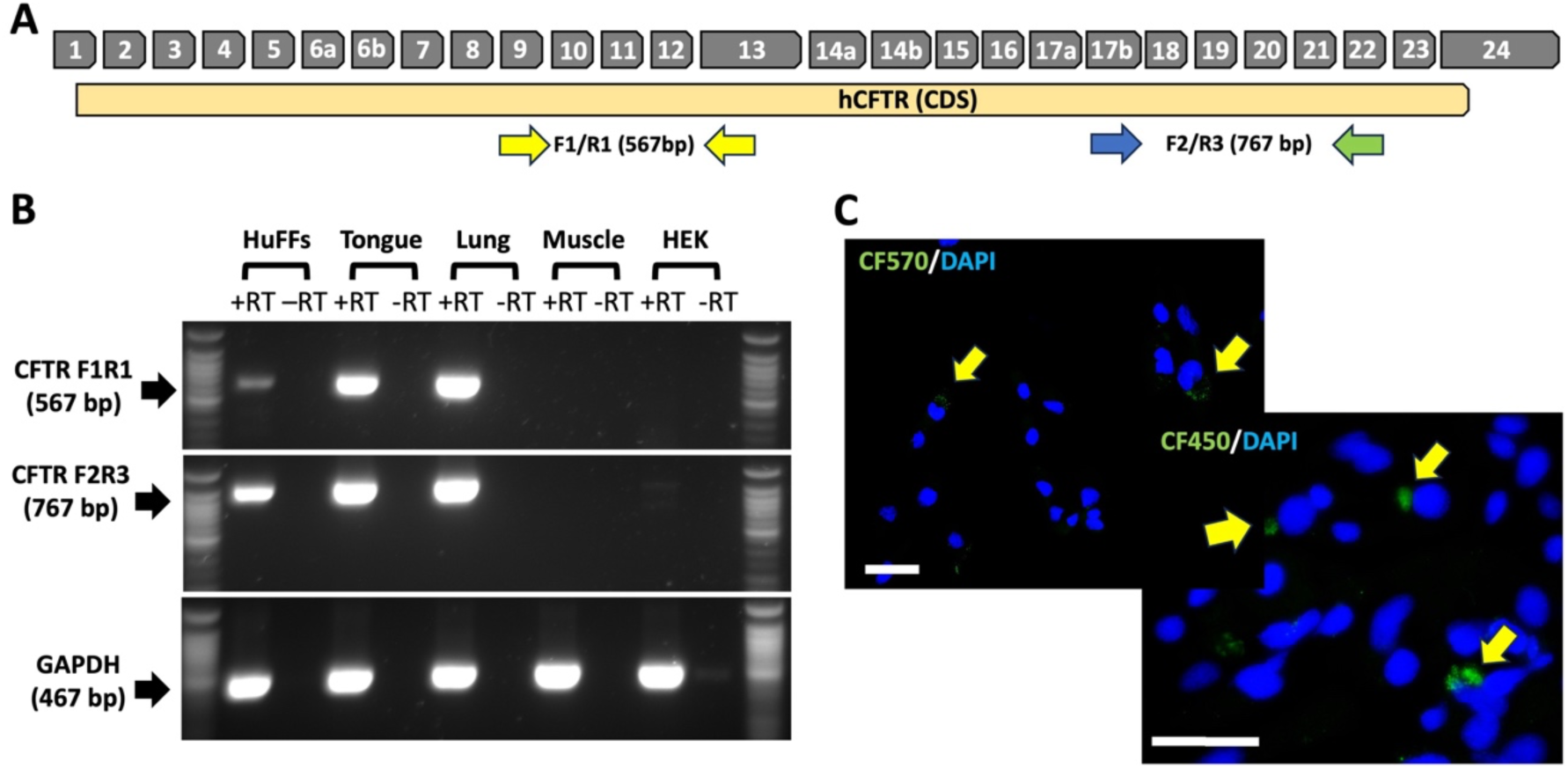
The CFTR transcript and protein are detected in human fungiform TBCs. **A-** Human CFTR gene structure, coding region, and location of the primers used to amplify a 567 bp fragment (F1/R1) across exon 9-13 encompassing nucleotide binding domain 1 and part of the regulatory domain, as well as a 767 bp fragment (F2/R3) spanning exon 17b-20 encoding a part of nucleotide binding domain 2. **B-** Agarose gel showing RT-PCR fragments amplified from human tongue, lung, HuFFs and HEK293 cells cDNA using CFTR-F1/R1, CFTR-F2/R3 or GAPDH primers. Note that the CFTR-F1/R1 fragment amplified from HuFFs is weak possibly uncovering alternative splicing of exon 9 in these cells [42]. NGS sequencing of the F1/R1 and F2/R3 PCR fragments from human tongue and HuFFs confirmed amplification of the CFTR without the ΔF508 mutation (data not shown). No reverse transcriptase (-RT) controls verify the absence of contamination. **C-** Immunocytochemical staining of HuFFs in culture showing punctate staining (Green – yellow arrow) of a subset of cells (∼ 5%) with two different mouse monoclonal antibodies (i.e. CF450 and CF570) directed against the regulatory domain of the human CFTR. Cell nuclei are stained with DAPI (Blue). Scale bar = 50 μm. No staining was observed when the primary antibody was omitted.

### CFTR channels expressed in HuFF cells impact the responsiveness of TBCs towards prototypical taste compounds

We assessed how the application of CFTR inhibitors might impact HuFF cell responsiveness to the bitter stimulus denatonium benzoate, a ‘fat’ stimulus linoleic acid, as well as to the sweetener sucralose. We selected denatonium and sucralose because CFTR function has been linked to G protein-coupled receptor-based taste receptor stimulation in other tissues [25, 26] and we selected linoleic acid hypothesizing that people with cystic fibrosis may be more sensitive to fatty acids because Orai1/ stromal interaction molecule 1 (STIM1) colocalization is increased in CF lung epithelial, interstitial, and luminal immune cells [27]. If also true in TBCs, this may affect taste responsiveness as STIM1 regulates calcium signaling in TBCs and preference for fat in mice [28].

**Table 1.** Sukumaran dataset (number of reads/ CV cell)

| Sukumaran dataset (number of reads/ CV cell) |  |  |  |  |  |  |  |  |  |  |  |  |  |
| --- | --- | --- | --- | --- | --- | --- | --- | --- | --- | --- | --- | --- | --- |
| Cell # | Type I |  |  | Type II |  |  |  |  | Type III |  |  |  |  |
|  | CFTR | ENTPD2 | SLC1A3 | PLCβ2 | TRPM5 | IP3R3 | Gg13 | Gust | SNAP25 | NCAM1 | CAR4 | PKD1L3 | GAD1 |
| Tas1 | 23 | 594 | 22 | 47,513 | 9,066 | 1,163 | 145 | 6 | 15 | 0 | 0 | 12 | 0 |
| Tas3 | 13 | 18 | 37 | 90,157 | 14,020 | 481 | 2,234 | 8 | 71 | 195 | 99 | 8 | 2 |
| Tas6 | 348 | 4530 | 28 | 5,130 | 1,724 | 836 | 103 | 1 | 66 | 2 | 0 | 165 | 0 |
| Tas8 | 2 | 785 | 39 | 15,844 | 6,189 | 2,161 | 91 | 61 | 41 | 4 | 0 | 5 | 2 |
| Tas9 | 2 | 179 | 64 | 14,044 | 4,788 | 1,137 | 35 | 2 | 78 | 29 | 0 | 6 | 8 |
| TIII9 | 1 | - | - | 6 | 0 | 1 | - | 1 | 118,985 | 5,304 | 5,265 | 12,290 | 14,146 |
| TIII10 | 874 | - | - | 3 | 1 | 2 | - | 6 | 119,582 | 847 | 9,226 | 19,838 | 57,430 |
| TIII13 | 5 | - | - | 0 | 1 | 0 | - | 0 | 95,218 | 14,715 | 1,029 | 4,797 | • 44,168 |

| Yamada dataset (CV and Fun) normalized data |  |  |  |  |  |  |  |  |  |  |  |  |  |
| --- | --- | --- | --- | --- | --- | --- | --- | --- | --- | --- | --- | --- | --- |
| Cell # | Type I |  |  | Type II |  |  |  |  | Type III |  |  |  |  |
|  | CFTR | ENTPD2 | SLC1A3 | PLCβ2 | TRPM5 | IP3R3 | Gg13 | Gust | SNAP25 | NCAM1 | CAR4 | PKD1L2 | GAD1 |
| CV 44480 | 3 | 1 | - | 8,197 | 5,599 | 1,365 | 16,016 | 61 | 1 | 4 | - | 3 | - |
| CV 519708 | 157 | 516 | - | 8,060 | 4,039 | 2,078 | 4,794 | 9,778 | 1 | 207 | - | 7 | - |
| CV 581198 | 1 | 43 | - | 1 | 0 | 0 | 0 | 0 | 0 | 0 | - | 0 | - |
| CV 754342 | 195 | 0 | - | 44 | 0 | 1 | 1 | 17 | 0 | 0 | - | 0 | - |
| Fun 63443 | 4 | 0 | 0 | 4,471 | 12 | 104 | 2,948 | 753 | 0 | 4 | - | 0 | - |

➤ 6.5% (4 of 62) of Gust-GFP CV cells express the CFTR at various levels (only cells that express CFTR are detailed in the table above). ➤ A subset 3.5% (1 of 29) Gust-GFP Fun cells express the CFTR at various levels. ➤ This data is in line with our IHC showing a small portion of mouse CV TBCs co-immunostained for Gust and CFTR as well as that of Merigo *et al.*

Calcium response amplitudes following denatonium exposure were significantly increased in the presence of the pharmacological CFTR inhibitor, CFTR-inh-172, compared to control stimulation in its absence (**Fig. 4A & B**; p = 0.003, Wilcoxon signed rank sum test). Sucralose evoked calcium amplitudes were also significantly larger with CFTR-inh-172 than without (**Fig. 4C**; p = 0.01). Calcium amplitudes following linoleic acid stimulation were not significantly affected by CFTR-inh-172 exposure (**Fig. 4D**). Although CFTR inhibition did not affect calcium amplitudes when HuFF cells were stimulated with linoleic acid, a significantly larger number of cells responded to linoleic acid when stimulated with CFTR-inh-172 compared to the number of these cells responding to linoleic acid alone (**Fig. 4E**; p = 0.002, McNemar’s chi square test with continuity correction). Interestingly, CFTR-inh-172 did not impact the number of cells responding to sucralose and denatonium.

**Figure 4:**
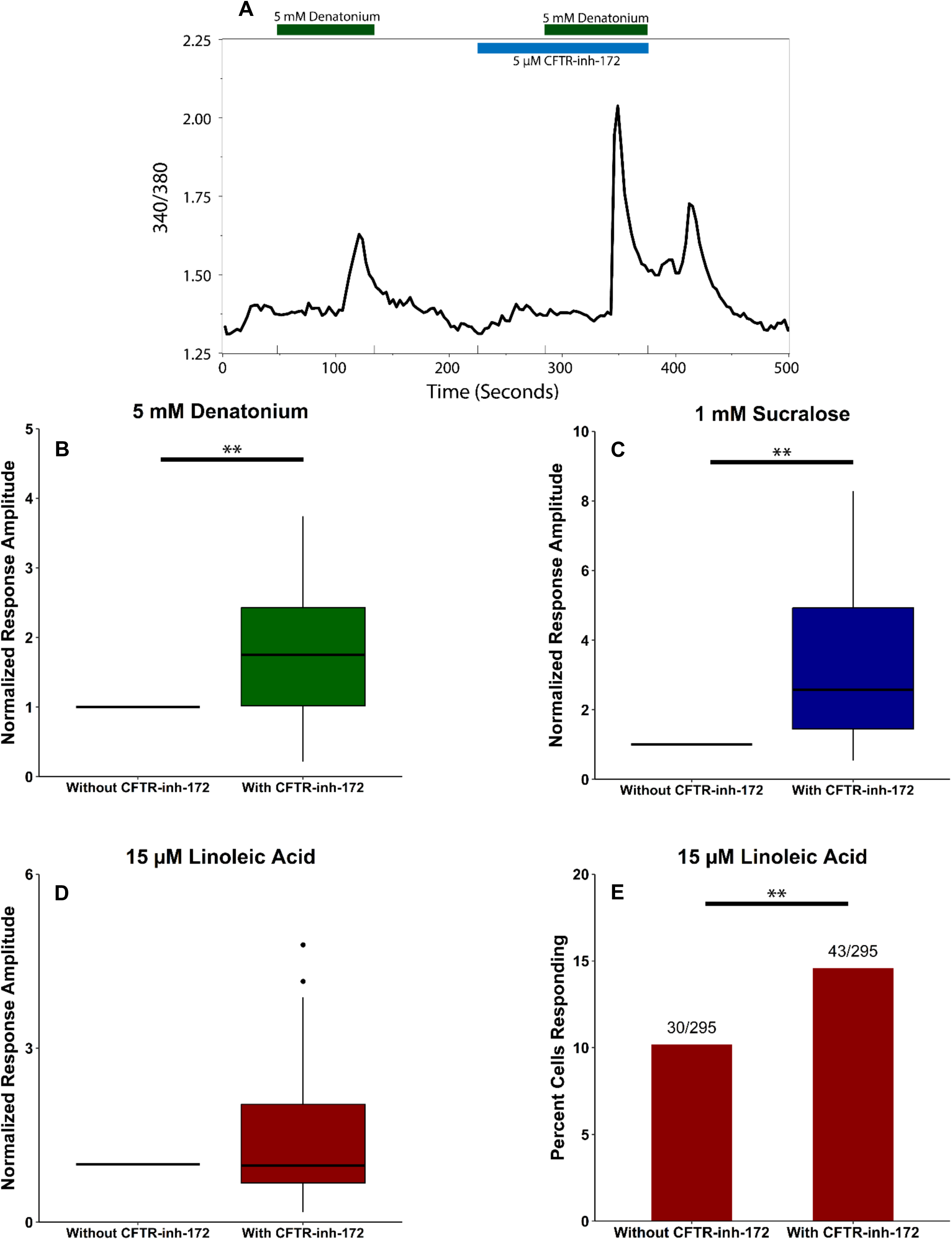
Acute CFTR inhibition modulates stimulus-evoked calcium responses in HuFF cells. **A**- A sample trace of a cell following the paradigm of exposure to a taste stimulus (e.g., denatonium) alone, followed by concurrent exposure with CFTR-inh-172. The 340/380 ratiometric fluorescence measure is plotted along the y axis, showing changes in cytosolic calcium. The bars on top of the graph, aligned with time on the axis, denote stimulus duration. **B**-Calcium response amplitudes following denatonium (5 mM) exposure were significantly increased in the presence of 5 μM CFTR-inh-172 compared to control stimulation in its absence (p = .003, Wilcoxon signed rank sum test, n = 19 cells). Responses in the presence of CFTR-inh-172 were normalized to responses to denatonium alone in corresponding cells. **C**-Sucralose (1 mM) evoked calcium amplitudes were significantly larger with CFTR-inh-172 than without (p = .01, n = 11 cells). **D**-Calcium amplitudes following linoleic acid stimulation were not significantly affected by CFTR-inh-172 exposure (p = .114, n = 30 cells). **E**-A significantly larger number of cells responded to 25 μM linoleic acid when concurrently stimulated with 5 μM CFTR-inh-172 compared to the number of the same cells responding to linoleic acid alone (30/295 vs. 43/295; p = .002, McNemar’s chi square test with Yates continuity correction).

Furthermore, incubation with CFTR-inh-172 (p = 0.007; data not shown) or another CFTR pharmacological inhibitor, Gly-H101, also enhances bitter-evoked ATP release from HuFF Cells (**Fig. 5**; p < 0.00001), suggesting that this modulation can be reflected throughout the signal transduction cascade.

**Figure 5:**
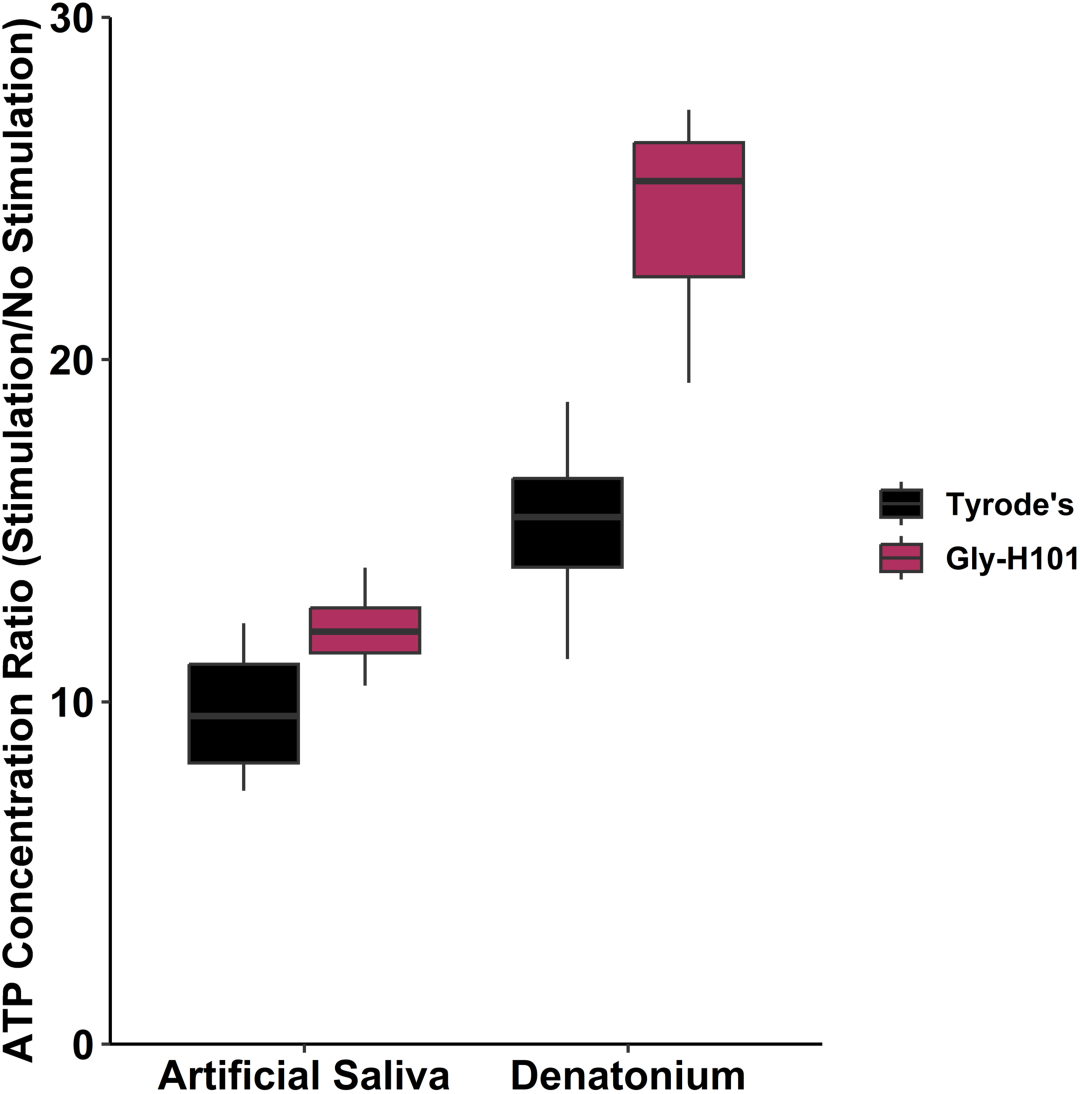
Incubation with CFTR inhibitors enhances bitter-evoked ATP release from HuFF cells. A significant (p < 0.00001) increase in extracellular ATP following bitter stimulation was observed after 30 min incubation with 43μM Gly-H101 relative to Tyrode’s buffer. No effect was observed with artificial saliva (p > 0.1; stimulus:incubation interaction: p < 0.01).

Preliminary analysis of HuFF cells infected with lentiviral CFTR shRNA (sc-108080 (CTRL) and sc-35054-V (CFTR), Santa Cruz) to knockdown CFTR indicate an impact of chronic CFTR inhibition on denatonium, sucralose, and linoleic acid evoked response amplitudes, as well as to linoleic acid evoked response rates (**Fig. 6**). At the time of this submission, CFTR knockdown experiments are still ongoing. Thus, while generally supporting the results of our calcium imaging and ATP secretion experiments, these CFTR knockdown experiments will need to be fully completed to assess significance and the extent to which the results from these experiments coincide with those generated using more acute methods of CFTR channel inhibition.

**Figure 6:**
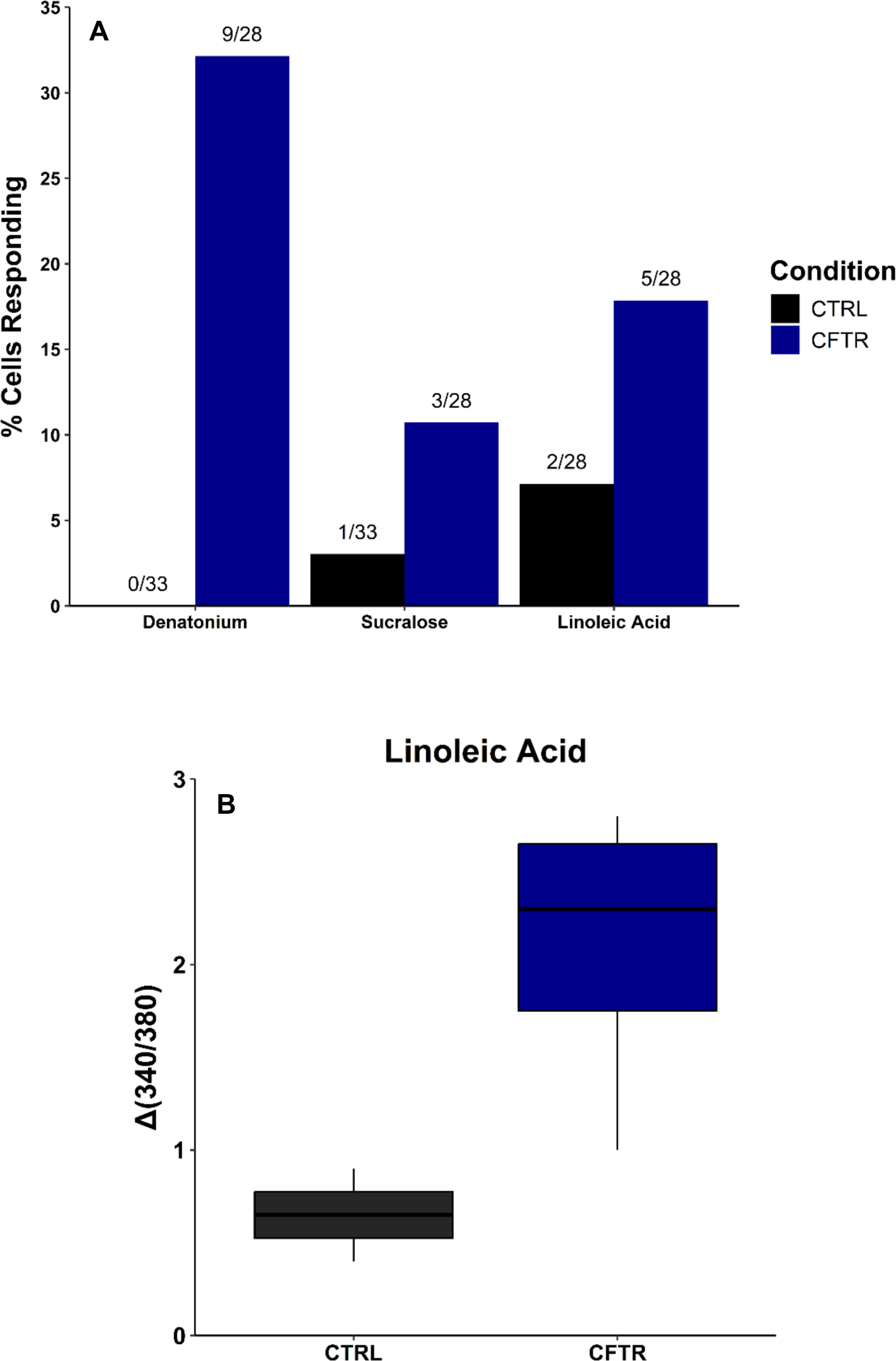
HuFF cells with lentiviral CFTR shRNA knockdown indicate an impact of chronic CFTR inhibition. **A-** A larger proportion of cells responded to 5 mM denatonium, 1 mM sucralose, and 25 μM linoleic acid after CFTR knockdown via lentiviral CFTR shRNA compared to the number of the same cells responding to these stimuli when the cells were infected with control shRNA. Exact response proportions are shown above each bar. **B-** Calcium response amplitudes following linoleic acid exposure appear increased following CFTR knockdown via lentiviral CFTR shRNA (n = 4 cells) compared to those of cells infected with control shRNA (n = 2 cells).

### HuFF cells produce CFTR-mediated chloride currents

Using an automated patch clamp system, we conducted voltage-clamp electrophysiological recordings of HuFF cells to determine the presence of chloride currents and whether CFTR contributes to these currents (**Fig. 7**). Application of forskolin, a cAMP stimulator and thus CFTR activator, increased the peak amplitude of outward chloride current, and this effect was reversed by the application of CFTR-inh-172 (**Fig. 7D**). These preliminary data therefore suggest that CFTR is active in HuFF cells.

**Figure 7:**
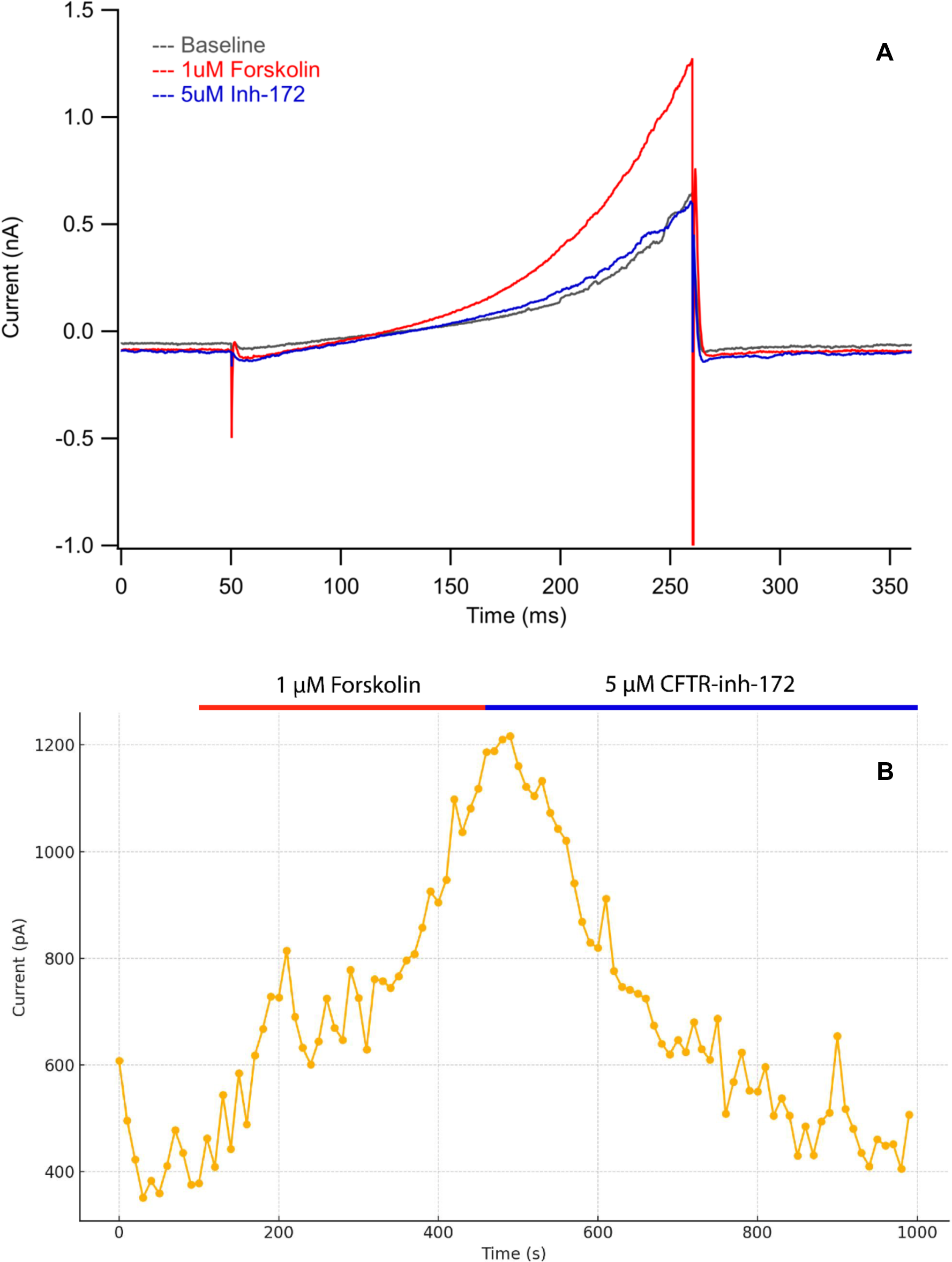
HuFF cells produce CFTR-mediated chloride currents. **A-** Representative trace of currents following forskolin treatment, measured before and after application of 5 μM CFTR-inh-172 (inh-172). **B-** Peak outward currents across several ramps over time with application of either forskolin (blue) or inh-172 (red) denoted by colored bars above the plot area. All data were collected from a single HuFF cell.

## DISCUSSION

The data presented in the current report demonstrate that CFTR is expressed both mouse and human TBCs and functionally active in human (TBCs. Using RT-PCR, immunocytochemistry, and electrophysiology, the study confirms the presence of CFTR transcripts, protein, and CFTR-mediated chloride currents in HuFF cells. Functional assays reveal that pharmacological inhibition of CFTR selectively enhances tastant-evoked signaling: bitter (denatonium) and sweet (sucralose) stimuli produce significantly larger calcium responses when CFTR is inhibited, whereas responses to the fatty acid linoleic acid show no change in amplitude but do show an increased proportion of responsive cells. CFTR inhibition also elevates bitter-evoked ATP release, suggesting amplification of downstream neurotransmitter or secondary effector signaling. Together, these findings indicate that CFTR modulates taste signaling in a tastant-specific manner and plays an active regulatory role in shaping peripheral taste transduction pathways.

### CFTR expression in other chemosensory cells

In addition to expression in TBCs, CFTR is also expressed in both murine and human nasal epithelial cells [29–31]. Interestingly, CFTR function has been linked to bitter taste receptor function in these cells. Bitter taste receptors (T2Rs) are expressed in airway cilia where they play a role in mucociliary clearance [32]. In addition to mediating cAMP-activated whole-cell Cl^-^ currents in nasal epithelial cells, CFTR function modulates bitter taste receptor mediated nitric oxide production and subsequent ciliary beat frequencies in these cells. It should be noted, however, that it remains unclear if Cl-conductance or changes in intracellular [Cl-] specifically affect T2R NO production [33]. It is interesting to speculate that CFTR function in nasal epithelial cells is coupled to T2R activation as stimulation of these receptors is known to activate phosphodiesterases that degrade cAMP (cyclic AMP).

### CFTR’s mechanism of action

CFTR functions as a channel that transports chloride and bicarbonate across epithelial surfaces. TBCs express an array of chloride currents, that display strong outward rectification, with both calcium– dependent and calcium–independent components [34]. For example, calcium-activated chloride channels such as TMEM16A (Anoctamin 1), are expressed in type 1 TBCs [35]. Stimulation of type 1 cells with ATP induces a TMEM16A-mediated current through the activation of metabotropic P2Y receptors [36]. The physiological roles of TMEM16A in TBCs remains unknown.

Based on our preliminary analysis, CFTR is thought to be expressed primarily in type 2 and 3 TBCs. CFTR acts as a gated ion channel, and its gating is regulated by two key factors: phosphorylation of the regulatory (R) domain and ATP binding and hydrolysis at two nucleotide-binding domains (NBDs) which are parts of CFTR [37, 38]. The R domain of CFTR contains multiple phosphorylation sites. Protein kinase A (PKA) is the primary kinase that phosphorylates these sites in response to elevated cAMP levels [37]. Phosphorylation is essential to relieve inhibition of the channel and allow it to open. Upon phosphorylation, ATP binds to both NBDs, promoting dimerization of the NBDs. This conformational change opens the channel pore [38].

CFTR is primarily activated by cAMP/PKA signaling, but calcium signaling also plays a major role in CFTR gating [39]. Patch-clamp studies have shown that calcium-loaded calmodulin can directly activate CFTR to levels comparable to PKA-mediated activation [40]. Indeed, calmodulin binding mimics PKA phosphorylation in promoting CFTR channel opening [40]. Interestingly, CFTR not only responds to Ca²⁺ but modulates calcium homeostasis by inhibiting ORAI calcium release-activated calcium modulator 1 (Orai1)-mediated Ca²⁺ entry and reducing endoplasmic reticulum Ca²⁺ release [39]. When certain taste receptors are activated by tastants, Ca²⁺ is released from internal stores. When CFTR signaling is blocked pharmacologically in TBCs, this release appears to be significantly augmented.

Taste modality-dependent differences in store-operated Ca^2+^ channel activation may account for the differential impacts of CFTR inhibition (i.e., response <u>enhancement</u> vs. <u>propensity</u> to respond). We speculate that the divergence of modulatory effects is mediated by Orai1 given that its utilization may be different between signal transduction mechanisms. For example, data from our lab suggests that denatonium and sucralose responses are dependent on ER calcium release while linoleic acid responses are not, though both depend on Orai1 (data not shown).

### Summary

Collectively, our data suggest that CFTR is functionally expressed in human TBCs and that its function can affect neurotransmitter release or secondary effector production, as has been observed in other tissues [25]. Indeed, continued characterization of CFTR’s functional influence on TBC signaling pathways will enhance our understanding of how tastants are transduced and how its function influences the output from the periphery to the central nervous system. Such knowledge could transform our understanding of taste transduction and uncover principles of cellular communication which would significantly advance the field and fill a longstanding gap in our knowledge. Moreover, TBCs share many characteristics with other cells of the gastrointestinal tract (e.g., pancreatic beta cells, enteroendocrine cells) [41]. Understanding the function of CFTR when expressed in these tissues may also provide some general insights into the functioning of this channel when expressed in other tissues.

