## Supplementary Tables for "The Cystic Fibrosis Transmembrane Conductance Regulator (CFTR) Modulates the Functional Output of Human Taste Bud Cells"

Table S1A: Primers used for RT-PCR experiments

| Gene | Forward primer | Reverse primer | | |
|---|---|---|---|---|
| GAPDH | ACCACAGTCCATGCCATCAC | ATGTAGGCCATGAGGTCCAC | NM_008084.4 | 467 |
| hCFTR (F1R1) | GACAGTTGTTGGCGGTTGC | AGTCTGGCTGTAGATTTTGGAGT | NM_000492.4 | 567 |
| hCFTR (F2R3) | CTGCGCTGGTTCCAAATGAG | TTGTGGCCATGGCTTAGGAC | NM_000492.4 | 767 |
| mCFTR (F0R0) | GCGATGCTTTTTCTGGAGATT | TCACTTGTAAAGGAGCAATCCATA | NM_021050.2 | 404 |
| mGLAST | GGTAAAATCGTGCAGGTCAC | CCACACCATTGTTCTCTTCC | NM_148938.3 | 673 |
| mTRPM5 | GTCTGGAATCACAGGCCAAC | GTTGATGTGCCCCAAAAACT | NM_020277.2 | 235 |
| mNCAM | GAACTCCATCAAGGTGAACC | CTGAACACAAAGTGAGCTGC | NM_010875.4 | 228 |

Table S1B: Assays used for qRT-PCR experiments

| Gene | Dye | TaqMan |
|---|---|---|
| GAPDH | VIC-MGB (Primer-limited) | Hs02786624_g1 |
| CFTR | FAM-MGB | Hs01565549_m1 |
